# A New Family of Similarity Measures for Scoring Confidence of Protein Interactions using Gene Ontology

**DOI:** 10.1101/459107

**Authors:** Madhusudan Paul, Ashish Anand

## Abstract

The large-scale protein-protein interaction (PPI) data has the potential to play a significant role in the endeavor of understanding cellular processes. However, the presence of a considerable fraction of false positives is a bottleneck in realizing this potential. There have been continuous efforts to utilize complementary resources for scoring confidence of PPIs in a manner that false positive interactions get a low confidence score. Gene Ontology (GO), a taxonomy of biological terms to represent the properties of gene products and their relations, has been widely used for this purpose. We utilize GO to introduce a new set of specificity measures: Relative Depth Specificity (RDS), Relative Node-based Specificity (RNS), and Relative Edge-based Specificity (RES), leading to a new family of similarity measures. We use these similarity measures to obtain a confidence score for each PPI. We evaluate the new measures using four different benchmarks. We show that all the three measures are quite effective. Notably, RNS and RES more effectively distinguish true PPIs from false positives than the existing alternatives. RES also shows a robust set-discriminating power and can be useful for protein functional clustering as well.

## 1 Introduction

A significant amount of protein-protein interaction (PPI) data has become available due to high-throughput technologies. PPI data play a central role towards a systems-level understanding of cellular processes with important applications in disease diagnosis and therapy. A considerable fraction of interactions are false positives due to limitations of experiments used in detecting protein interactions [1]. Hence, a ranking or a scoring mechanism distinguishing between true and false interactions is important for any downstream analysis. There have been continuous efforts to utilize additional knowledge resources, such as Gene Ontology (GO) [2], in scoring confidence of PPIs in a manner that false positive interactions get a low confidence score [3]. The primary objective of this work is to introduce a new family of semantic similarity measures (SSMs) between gene products using GO for scoring confidence of PPIs.

GO has been effectively utilized in predicting and validating PPIs [4], [5], [6], and confidence scoring of PPIs [7], [8], [9], [10], [11], [12] among other genomic applications such as predicting protein functions [13], [14], [15], analyzing pathways [16] etc. It is a taxonomy of biological terms to represent the properties of genes and gene products (e.g., proteins) and is organized as a directed acyclic graph (DAG) to describe the relationship among the terms. GO is made up of three independent ontologies: biological process (BP), cellular component (CC), and molecular function (MF). A section of GO DAG (Release March 2015) is shown in the Supplementary Material. Terms closer to the root are more generic in nature and specificity of terms gradually increases as we move towards the leaves. The more specific a term is, the more informative it is. Ontology-based SSM is a quantitative function that measures the similarity between two terms based upon their relations over a set of terms organized as an ontology. Formally, it is a function of two ontology terms (or two sets of ontology terms) that returns a real number indicating the closeness between the terms in the context of semantic meaning [3]. Gene or gene products in different model organisms are annotated to GO terms based on various evidences and is available through annotation corpora. An annotation corpus of a species (e.g., yeast) is an association between gene products of the species and GO terms.

### 1.1 Motivation and Hypothesis

The notion of Information Content (IC) is widely used in defining SSMs. It quantifies specificity of a term in an ontology, i.e., how specific a term in an ontology is. The IC is explained formally in section 2. The IC based SSMs assume that the given ontology is complete and define specificity of a term by considering the whole ontology. However, GO is being updated regularly with the addition of new terms and removal of old terms. Furthermore, when new information of gene or gene product is discovered, annotation data corresponding to the appropriate terms are updated as well. Some proteins are annotated with a large number of terms, while many proteins are annotated to one term only, i.e., annotations are not uniformly distributed among the terms (annotation bias). Thus the continuous evolution of the GO DAG, regular updates in annotation and nonuniform distribution of terms (as well as annotations) over the ontology are likely to impact confidence scores of several PPIs with each update.

A GO term is more closely related to its ancestors and descendants as the ontology is hierarchically organized. The major part of contribution towards specificity of a term is accumulated through the properties of its ancestors and descendants. Therefore for quantifying specificity of a term in an ontology like GO (which is very large, complex, continuously evolving and not uniformly distributed), it is safe to consider the properties of the subgraph consisting of the term itself along with its ancestors and descendants only instead of considering the whole ontology, to minimize the impact of continuous evolution.

Our main hypothesis is that the explicit encoding of the aforementioned unexplored subgraph-based specificity notions into a new family of SSMs could be useful for scoring confidence of PPIs.

### 1.2 Definition of the Problem and Contribution

The main problem of the current study is to define the specificity of a GO term, based on the properties of the subgraph consisting of the term itself along with its ancestors and descendants only, that could be useful for scoring confidence of PPIs.

With the aforementioned unexplored notion of specificity, we introduce three simple yet effective specificity measures: Relative Depth Specificity (RDS), Relative Node-based Specificity (RNS), and Relative Edge-based Specificity (RES). This new set of specificity measures led to a new family of SSMs.

We compare the performance of the new SSMs with six state-of-the-art SSMs proposed by Resnik [17], Lin [18], Schlicker *et al.* [19], Jiang & Conrath [20], Wang *et al.* [21], and Jain & Bader [22], referred to as *Resnik, Lin, Rel, Jiang, Wang*, and *TCSS*, respectively, in the rest of the paper. Resnik and TCSS have been considered to be the best SSMs for scoring confidence of PPIs by several studies such as Guo *et al.* [23], Xu *et al.* [24], Jain & Bader [22], and Pesquita [25]. We use four different benchmarks to evaluate the new SSMs. The four benchmarks are: 1) correlation with reference dataset from HIPPIE database [26], 2) ROC curve analysis with DIP database [27], 3) set-discriminating power of KEGG pathways [28], and 4) correlation with protein family (Pfam) using CESSM dataset [29]. The first benchmark is for human PPIs only as HIPPIE is an integrated database of human PPIs and the rest of the three benchmarks are applied to both yeast (S. cerevisiae) and human (H. sapiens) PPIs.

The rest of the paper is organized in the following manner. A brief survey of the literature is presented in section 2. The new family of SSMs is explained in section 3. Section 4 describes the experimental design, evaluation metrics, datasets used, implementation and results. In section 5, results are analyzed and discussed. Finally, section 6 introduces the conclusions and future work.

## 2 Related Work

This section introduces a brief review of the literature on PPI confidence scoring methods and GO-based SSMs. For an indepth review of the family of GO-based SSMs, we refer the reader to the surveys by Pesquita *et al.* [3], Harispe *et al.* [30], Mazandu *et al.* [31], and Pesquita *et al.* [25].

### 2.1 PPI Confidence Scoring Methods

Computational approaches for scoring confidence of PPIs mainly differ in the selection of information used in the prediction model. The common sources of this information are three-dimensional protein structures [32], protein sequences [33], gene expression profiles [34], phylogenetic trees [35], [36], phylogenetic profiles [37], GO [7], [8], [9], [10], [11], [12] etc. Some approaches utilize topology of interaction network from already existing true PPIs [38], [39], [40]. Text mining on peer-reviewed literature is also used for scoring confidence of PPIs [41]. A few approaches utilize multiple sources of information [42], [43]. However, GO is a very comprehensive resource for the properties of gene products and their functional relationships across species. It provides a promising way to infer functional information of gene products. The idea of semantic similarity is a common way to utilize GO for scoring confidence of PPIs. Semantic similarity between two proteins (see section 2.3) involved in a PPI may be treated as a confidence score of the interaction. The current study is primarily focused on the SSMs by exploiting GO for scoring confidence of PPIs.

### 2.2 Go-based SSMs

Ontology-based SSMs were originally introduced in the fields of cognitive sciences by Tversky [44] and Natural Language Processing (NLP) and Information Retrieval (IR) by Rada *et al.* [45]. Since then a plethora of semantic similarity measures based on WordNet (a large lexical database of English) were developed such as the pioneering works introduced by Resnik [17], Jiang & Conrath [20] and Lin [18]. However, the first pioneering work was introduced by Lord et al. [46], [47] in the field of biology and this work has started the research on the development of GO-based SSMs and their applications in genomics such as [19], [21], [22], [48], [49], [50]. Here, we provide a brief overview of different SSMs.

Existing SSMs are classified broadly into two categories: *edge- and node-based* [3]. Edge-based measures are the natural and direct way of defining SSMs. Rada *et al.* [45] introduced a SSM of this kind in a lexical taxonomy, which was then applied in GO by Nagar and Al-Mubaid [48]. Subsequently, several edge-based SSMs have been developed and used in GO [51], [52], [53], [54]. In the edge-based approach, shared paths between two terms are primarily considered to compute the similarity between them. It assumes that terms at the same level have similar specificity and edges at the same level represent same semantic distances between two terms [3], which are seldom true in GO. Furthermore, an edge-based approach does not account annotation information of terms and entirely relies on the topological structure of the GO DAG. Hence edge-based methods are more sensitive to the intrinsic structure of the GO DAG.

The most commonly used SSMs are node-based that compute the similarity between two terms by comparing their properties, common ancestors, or their descendants. As mentioned earlier, majority of the node-based approaches use the notion of information content (IC) to define the specificity of a term. The IC of a term *t* is defined as

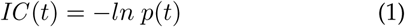

where *p*(*t*) is the probability or frequency of occurrence of *t*. Usually, the descendants of *t* are also considered for computing *IC*(*t*). The probability of occurrence, *p*(*t*) of term *t* in GO is defined as:

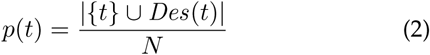

where *Des* (*t*) is the set of descendants of *t* and *N* is the number of terms in the ontology. Since gene products are annotated to terms in GO, *p*(*t*) is estimated as the frequency of annotations of *t*, i.e.,

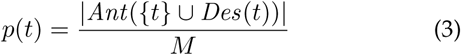

where *Ant*(*T*) is the set of annotations to the set of terms *T* and *M* is the total number of annotations in the GO. In words, it is the ratio of the number of annotations to *t* and its descendants to the total number of annotations. The aforementioned two definitions are commonly known as an *intrinsic* and *extrinsic* way of defining the probability function *p*(*t*), respectively.

The most commonly used node-based SSMs are Resnik [17], Lin [18], and Jiang & Conrath [20], which were initially developed for WordNet and subsequently applied to GO by Lord et *al.* [46], [47]. Thereafter, a number of node-based SSMs have been proposed in order to improve the existing SSMs in different perspectives and applications [19], [55], [56], [57], [58], [59], [60], [61], [62], [63], [64], [65], [66]. The major drawbacks of IC based SSMs are already pointed out in section 1.1. SSMs such as [21], [50], [67], [68] combine both node- and edge-based approaches and are commonly referred to as hybrid approaches. Recently, some complex structural-based SSMs are also developed [22], [69], [70].

### 2.3 SSM between Two Sets of Terms

A gene product may be annotated with more than one term in the same GO. Suppose, *p*_1_ and *p*_2_ are two gene products annotated to the set of terms *S* and *T*, respectively. The similarity between *p*_1_ and *p*_2_ are calculated as the similarity between two sets *S* and *T*, i.e., SSM(*p*_1_,*p*_2_) = SSM(*S*, *T*). Therefore we need to combine GO terms of S and T. Generally, the following three types of strategies used in the literature:

#### Maximum (MAX)

In MAX strategy [71], similarity between *S* and *T* is calculated as the maximum of the set *S* × *T*.

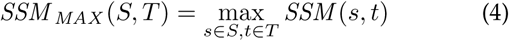

#### Average (Avg)

In ‘average’ strategy [46], [47], similarity between *S* and *T* is calculated as the average of the set *S* × *T*.

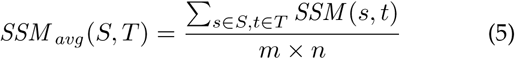

where *m* = |*S*| and *n* = |*T*|.

#### Best-match average (BMA)

SSMs between two sets of terms form a matrix. BMA [19], [72], [73] is defined as the average of all maximum SSMs on each row and column of the matrix.

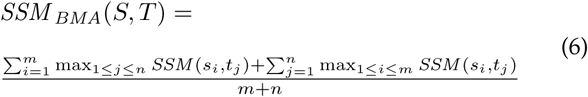

where *s_i_* ∈ *S* and *t_j_* ∈ *T*.

### 2.4 SSMs used in Evaluation

#### Resnik

Resnik considers IC of the *most informative common ancestor* (MICA) only [17]. The similarity between two terms *s* and *t* in Resnik is defined as

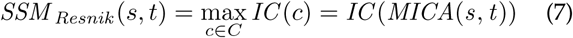

where *C* is the set of common ancestors of *s* and *t*, and IC is the information content defined earlier. It is the IC of the closest common ancestor or *lowest common ancestor* (LCA) of s and t.

#### Lin and Jiang

Although Resnik is very effective for computing information shared by two terms, it cannot distinguish between pairs of terms having the same MICA. To overcome the problem, Lin and Jiang are developed by considering ICs of both the terms along with their MICAs in different ways [18], [20]. The similarity between two terms is calculated by these two methods as

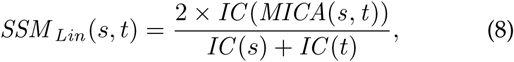

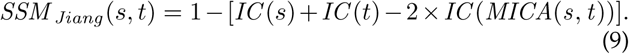

#### Rel

Lin and Jiang overestimate when one term is an ancestor of another. For example, when both the terms are same, the similarity score will be *1*, irrespective of its specificity. Rel combines Resnik and Lin in order to capture relevance information by multiplying one minus the *extrinsic* probability of MICA to *SSM_Lin_* [19]. As per Rel, the similarity between two terms is calculated as

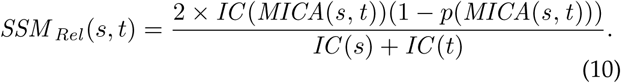

#### Wang

Wang is a hybrid measure that combines both edge- and node-based approaches [21]. Let *G_t_* = (*V_t_*, *E_t_*) be a DAG for a term t in GO such that *V_t_* is the set of ancestors of *t* including *t* itself and *E_t_* is the set of edges connecting terms in *G_t_*. Terms closer to term *t* in *G_t_* contribute more of its semantics to the semantics of term *t*. The semantic contribution of a term *c* to the semantics of term *t* in *G_t_* is denoted as S-value of *c* or *S_G_t__* (*c*) and defined as:

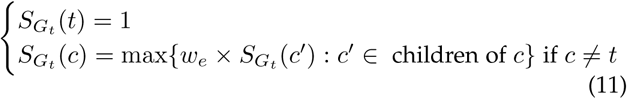

where *w_e_* (0 < *w_e_* < 1) is semantic contribution factor for edge *e* ∈ *E_t_* from term *C* to term *c*. For example, semantic contribution factors (*w_e_*) of *is_a* and *part_of* relationships may be treated as 0.8 and 0.6, respectively. To compare semantics of two terms, a semantic value *SV*(*t*) is computed as the aggregate contribution of the semantics of all the terms in *G_t_* to term *t* and defined as:

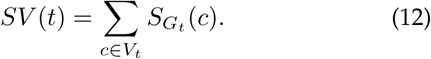

Now, SSM between two terms *s* and *t* with respect to their DAGs *G_s_* = (*V_s_*, *E_s_*) and *G_t_* = (*V_t_*, *E_t_*) is defined as:

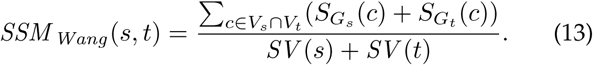

The numerator is the summation of S-values of common terms between the two DAGs. S-values of common terms between the two DAGs may not be same as the locations of *s* and *t* may differ in GO.

#### TCSS

TCSS exploits the unequal depth of biological knowledge representation in different branches of GO DAG [22]. The objective of TCSS is to identify subsets of similar GO terms (e.g., terms related to nucleus and terms related to mitochondrion belong two different subsets) and score PPIs higher if participating proteins belong to the same subset compared to PPIs whose participating proteins belong to different subsets. The authors have introduced a structural-based IC, referred to as topological information content (ICT), to identify sub-graph root terms during preprocessing stage.

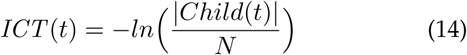

where *Child*(*t*) is the set of children of *t* and *N* is the number of terms in the ontology.

## 3 The New GO-based SSMs

In this section, we introduce the new family of SSMs based on the proposed set of specificity measures. To define specificity of a GO term we consider the properties of the subgraph consisting of the term itself along with its ancestors and descendants only and ignore the rest of the ontology. The new specificity models quantify how specific a term in ontology is. The specificity of a parent (term) always will be less than any of its children. RDS considers a specific path of the aforementioned subgraph, while RNS and RES consider the whole subgraph. However, RNS relies on the properties of the nodes only, whereas RES considers the edges of the subgraph as well.

### 3.1 Relative Depth Specificity (RDS)

Let *d_t,r_* and *d_l,t,r_* are length of the longest path from term *t* to the root *r* and length of the longest path from any leaf *l* to the root *r* via the term *t*, respectively. Then, RDS of a term *t* in GO is defined as

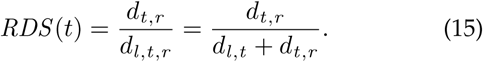

In words, *RDS* (*t*) is the ratio between the length of the longest path from the term *t* to the root and the length of the longest path from any leaf to the root via the term *t*. This is the simplest SSM that does not consider annotation information. The specificity of the leaves and the root would be highest (1) and lowest (0), respectively. When multiple paths are present between two terms, we consider the longest one as it is likely to be more informative than others.

### 3.2 Relative Node-based Specificity (RNS)

Let *G*_1_ (*V*_2_, *E*_2_) be the subgraph consisting of the term *t* itself along with its ancestors; and *G*_2_(*V*_2_,*E*_2_) be the subgraph consisting of the term *t* itself along with its ancestors and descendants. The RNS of a term *t* in GO is defined as

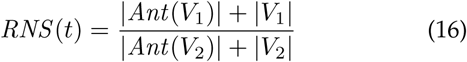

where *Ant*(*T*) be the set of annotations to the set of terms *T* as mentioned earlier. In words, it is the ratio of the sum of nodes along with its annotations of the subgraph consisting of the term *t* and its ancestors to the sum of nodes along with its annotations of the subgraph consisting of *t*, its ancestors and descendants. Thus, RNS of the leaves and the root would be highest (1), and lowest (close to 0), respectively.

### 3.3 Relative Edge-based Specificity (RES)

We define the weight of an edge *e*(*t*_1_, *t*_2_) between terms *t*_1_ and *t*_2_ in GO as:

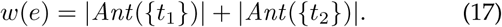

It is the summation of the number of annotations of terms *t*_1_ and *t*_2_. The weight of a set of edges *E* is defined as:

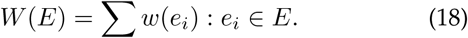

It is the summation of weights of all edges in the set of edges *E*. Let *G*_2_(*V*_2_, *E*_2_) be the subgraph consisting of the term *t* itself along with its ancestors and *G*_2_(*V*_2_,*E*_2_) be the subgraph consisting of the term *t* itself along with its ancestors and descendants as in RNS. The Relative Edge-based Specificity of a term *t* in GO is defined as

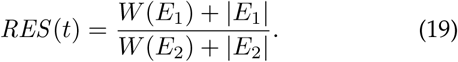

In words, it is the ratio of the summation of weighted and unweighted edges of the subgraph consisting of the term *t* itself along with its ancestors to the summation of weighted and unweighted edges of the subgraph consisting of *t* itself along with its ancestors and descendants. Thus, specificity of the leaves and the root would be highest (1), and lowest (0), respectively.

The similarities between the two terms *s* and *t* are calculated as:

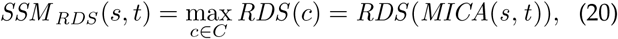

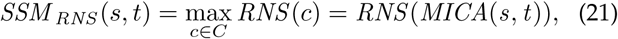

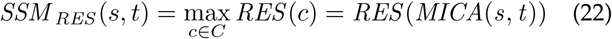

where *C* is the set of common ancestors of *s* and *t* as mentioned earlier.

We have chosen the MICA to define the shared specificity between the two terms similar to Resnik. It is noteworthy to mention that the proposed specificity models are different from IC models as they do not rely on probability functions. Therefore we cannot directly apply the new specificity models to other IC-based similarity measures such as Lin, Rel, and Jiang.

## 4 Evaluation

In this section, we detail the experimental design, evaluation metrics, datasets used, implementation and results. As already mentioned, six state-of-the-art SSMs are chosen as baseline methods and four benchmarks are considered for evaluation of the new SSMs.

### 4.1 Experimental Setup

Our experimental design for evaluation is based on the following two assumptions. First, two proteins involved in the same biological process(es) are more likely to interact than proteins involved in different processes [5, p.953] and [22]. Second, two proteins need to come in close proximity (at least transiently) for interaction, hence co-localization also provides evidence of interaction [74, p. 689] and [22]. However, if two proteins interact physically, there is no guarantee that they share the same molecular function [75, p. 27]. The ‘average’ strategy underestimates when two gene products share many similar terms as it considers all possible term pairs of the two gene products [76]. By contrast, the MAX strategy overestimates when two gene products share few similar terms as it is indifferent to the number of dissimilar terms between the gene products [76]. The BMA strategy, which considers both similar and dissimilar terms [76], does not suffer from the aforementioned limitations. Further, in PPIs, proteins need to be in close proximity (share similar CC terms) and participate in the same biological process (share similar BP terms) once, among all possible combinations, to become biologically relevant [22]. Hence MAX and BMA are considered better strategies than the ‘average’ for scoring confidence of PPIs. In light of the above discussion, we use BP and CC ontologies of GO along with MAX and BMA strategies for performance evaluation. We exclude electronically inferred annotations (IEA) of GO terms which lack manual curation. We consider only those protein pairs which are having both the proteins annotated with at least one GO term other than the root in their respective ontologies.

As mentioned earlier, the new SSMs are evaluated on the four benchmarks: 1) correlation with reference dataset from HIPPIE database, 2) ROC curve analysis in predicting true PPIs from DIP database, 3) set-discriminating power of KEGG pathways, and 4) correlation with Pfam on CESSM dataset. Evaluation is done using both yeast (S. cerevisiae) and human PPIs except for the first benchmark, as it contains only human PPIs. We use Entrez and ORF gene ids for human and yeast, respectively, except while comparing with TCSS where UniProtKB and SGD gene ids were used for human and yeast, respectively. We have not performed the comparison with TCSS on the second and third benchmarks (for human) as some UniProtKB ids (after mapping from Entrez ids) were not found in the annotation.

### 4.2 Evaluation Metrics and Baselines

This section introduces how and why each benchmark is used for evaluation. A brief outline and formulation of each metrics used are presented here.

#### 4.2.1 Correlation with Reference Dataset from HIPPIE Database

The HIPPIE database [26] integrates most of the publicly available PPI databases like BioGRID [77], DIP [78], HPRDS [79], IntAct [80], MINT [81], BIND [82], MIPS [83]. It also includes interactions from several manually selected studies. The HIPPIE score of a PPI is defined by considering the following parameters: the number of studies where the PPI was detected, the number and quality of the experimental techniques used to detect the PPI, and the number of non-human organisms where the PPI was reproduced. The authors of HIPPIE showed that their scoring scheme of interactions correlates with the quality of the experimental characterization. We use a reference dataset from HIPPIE database to evaluate different SSMs. Pearson correlation is calculated between the HIPPIE score and PPI confidence score obtained using an SSM.

#### 4.2.2 ROC Curve Analysis

Similarity measures can be treated as binary classifiers to classify a given PPI as positive or negative with a reasonable cutoff similarity score. PPIs having similarity score greater than the cutoff are treated as positive. Receiver operating characteristic (ROC) curve analysis is used to evaluate the performance of a binary classifier. ROC curve is a graph plotting of true positive rate (TPR or sensitivity) against false positive rate (FPR or 1-specificity) by varying discrimination threshold or cutoff. The area under the ROC curve (AUC) is the measure of discrimination, i.e., the ability of the classifier to classify correctly. An AUC of 1 represents perfect classifier. We utilize the core subsets of yeast and human PPIs from the DIP database [27] to evaluate different SSMs for AUCs.

#### 4.2.3 Set-discriminating Power of KEGG Pathways

A biological pathway is a sequence of biochemical steps to accomplish a specific biological process within a cell. Therefore proteins involved in a pathway are more likely to interact among themselves than the proteins belonging to different pathways. Proteins within a pathway are likely to be annotated with the same or similar terms in GO too and should show high similarity scores. We consider three sets of selected KEGG pathways [28] to evaluate different SSMs for their discriminating power as discussed in the following paragraph.

For each KEGG pathway, an *intra-set average similarity* is calculated as the average of all pairwise similarities of proteins within the pathway. An *inter-set average similarity* for every two pathways is also calculated as the average of all pairwise SSMs of proteins between the two pathways. During calculation of *inter-set average similarity*, we do not consider those pairs whose both the proteins are same. A discriminating power (DP) of a pathway is defined in [84] as the ratio between *intra-set average similarity* and the average of all *inter-set average similarities* between the chosen pathway and rest other pathways. Let *P* = {*P*_1_, *P*_2_,…, *P_n_*} be the set of KEGG pathways, each pathway *P_k_* contains *m_k_* number of proteins and *p_ki_* denotes *i^th^* protein in *P_k_*.

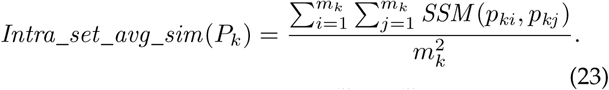

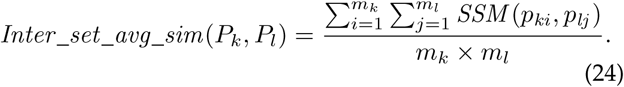

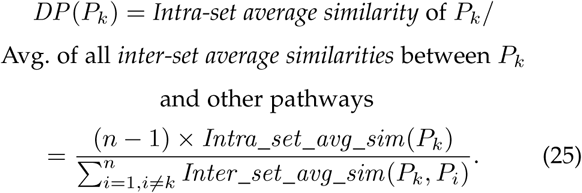

#### 4.2.4 Correlation with Protein Family (Pfam)

A protein family (Pfam) is a group of proteins that are evolutionarily related, i.e., they share a common evolutionary ancestor. Proteins belonging to a family often show functional similarity. The Jaccard index is used to calculate Pfam similarity. The Jaccard index of two proteins is calculated as the ratio of the number of protein families they share to the total number of protein families they belong. We utilize dataset of protein pairs used in CESSM [29]. For each pair, Pfam similarity (Jaccard index) and similarity scores of different SSMs are calculated and finally, the Pearson correlation between the two scores is obtained.

### 4.3 Datasets

In this section, we describe the sources of different datasets used in the evaluation and the corresponding preprocessing steps. A summary of the datasets used is presented in Table 1.

**TABLE 1.**
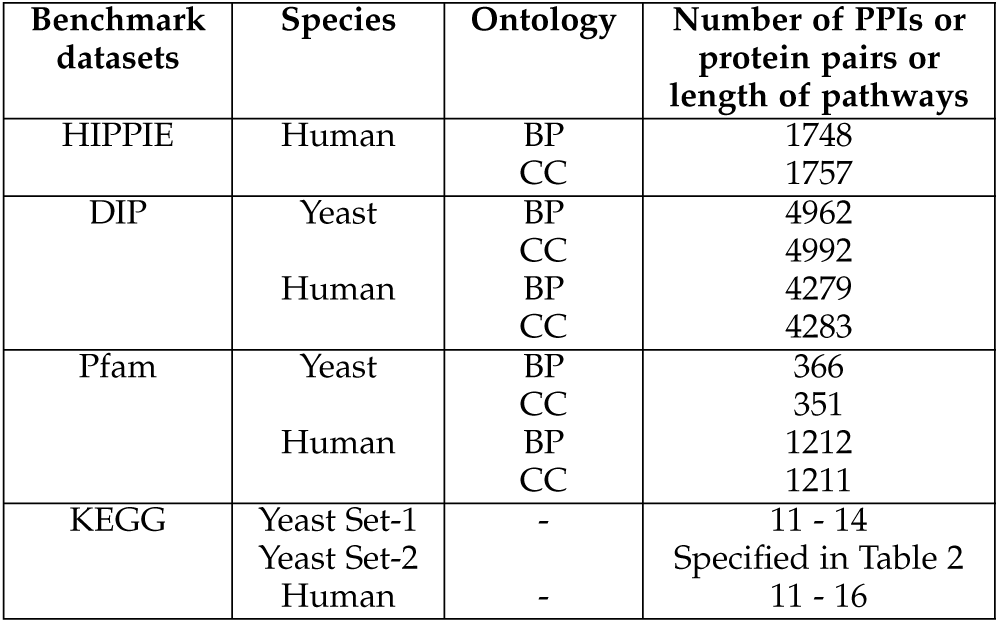
Summary of Datasets used in Evaluation

#### 4.3.1 Reference Dataset from HIPPIE Database

We download Human Integrated Protein-Protein Interaction rEference (HIPPIE) dataset on 09.01.2015 [26]. We extract one reference dataset from HIPPIE consisting of PPIs detected by four top-scored experimental techniques: far-Western blotting, isothermal titration calorimetry, nuclear magnetic resonance, and surface plasmon resonance experiments as in [85]. The interaction detected by any of the chosen four experimental techniques have a high probability of being an actual interaction [85]. The number of PPIs present in the reference datasets is shown in Table 1.

#### 4.3.2 Datasets for ROC Curve Analysis

We download the core subsets of PPIs from the Database of Interacting Proteins (DIP) [27] for S.cerevisiae and H.sapiens on 29.10.2015. DIP is a database of experimentally detected PPIs from various sources. We assume that these interactions are real and treat them as positive instances of interactions. DIP uses UniProt Ids for proteins. We map UniProt Ids into Entrez and ORF gene Ids for human and yeast, respectively. Table 1 shows the number of PPIs of DIP dataset used in this study. As done in [22], an equal number of negative PPI datasets are independently generated by randomly choosing protein pairs annotated in BP and CC, and are not present in the iRefWeb database [86] (version date: 27.11.2015), a combined database of all known PPIs.

#### 4.3.3 KEGG Pathways

We extract two sets of KEGG pathways [28] of each of the two organisms, S.cerevisiae and H.sapiens, using org.Sc.sgd.db and org.Hs.eg.db packages with R 3.1.2 version. The first set contains a number of genes between 11 to 14 and the second set 11 to 16. We choose the above ranges so that each set contains the same (11) number of pathways and takes a reasonable time to compute. The two sets have three common pathways: Terpenoid backbone biosynthesis (sec00900 and hsa00900), Riboflavin metabolism (sec00740 and hsa00740), and Pantothenate and CoA biosynthesis (sec00770 and hsa00770). However, each of them is from different organisms and may not show similar results. Another set of 11 yeast KEGG pathways (Table 2) with more diverse functionality is also considered to get a broader insight into the inter-set discriminating power.

**TABLE 2.** List of 11 Yeast Pathways with More Diverse Functionality used in the Study.

| Category | Subcategory | Pathway Id | Pathway Name | No. of Genes |
| --- | --- | --- | --- | --- |
| Metabolism | Carbohydrate metabolism | sce00040 | Pentose and glucuronate interconversions | 10 |
|  | Energy metabolism | sec00920 | Sulfur metabolism | 15 |
|  | Lipid metabolism | sec00565 | Ether lipid metabolism | 5 |
|  | Amino acid metabolism | sec00360 | Phenylalanine metabolism | 9 |
|  | Glycan biosynthesis and metabolism | sec00514 | Other types of O-glycan biosynthesis | 13 |
|  | Metabolism of cofactors and vitamins | sec00750 | Vitamin B6 metabolism | 11 |
|  | Metabolism of terpenoids and polyketides | sec00900 | Terpenoid backbone biosynthesis | 13 |
|  | Metabolism of other amino acids | sec00410 | beta-Alanine metabolism | 8 |
| Genetic Information Processing | Folding, sorting and degradation | sec04122 | Sulfur relay system | 8 |
|  | Replication and repair | sec03450 | Non-homologous end-joining | 10 |
| Environmental Information Processing | Signal transduction | sec04070 | Phosphatidylinositol signaling system | 15 |

#### 4.3.4 CESSM Dataset for Correlation with Pfam

The Collaborative Evaluation of GO-based Semantic Similarity Measures (CESSM) is an online tool for evaluation of GO-based SSMs against sequence, Pfam and EC similarities [29]. Since CESSM has been published around ten years ago, it uses ten years old dataset (August 2008 GO and GOA-UniProt). In the meanwhile, GO DAG, its annotation, as well as Pfam have substantially changed. Moreover, we use GO.db (version:3.1.2) and org.Hs.eg.db (version:3.1.2) R packages that utilize March 2015 GO and annotations, respectively, in the evaluation. Hence we could not use CESSM automated tool. However, we utilize the dataset of protein pairs used in CESSM to find correlation against Pfam similarity only, since GO captures the functional aspect of gene or gene products primarily. All pairs of proteins are mapped into Entrez and ORF gene Ids for human and yeast, respectively. The dataset involves 13,430 protein pairs of 1,039 proteins from various species. The authors of CESSM reported that both proteins of each pair are manually annotated to at least one term within all the three GOs with a uniform IC of at least 0.5 and have at least one EC class and one Pfam class. The number of protein pairs used for this evaluation is shown in Table 1.

### 4.4 Implementation

The new SSMs are implemented in the R programming language [87]. We use GOSemSim R package (version: 1.26.0) [88] for implementations of Resnik, Lin, Rel, Jiang, and Wang SSMs. For GO and corresponding annotations, we use GO.db, org.Sc.sgd.db (for yeast), and org.Hs.eg.db (for human) R packages (version:3.1.2, March, 2015 release) [89], [90], [91]. We maintain versions of all R packages so that they use same GO and corresponding annotations. For TCSS, we use the implementation provided by the authors with the default set of parameters. The original implementation of TCSS uses MAX strategy only. Therefore we modify it to include BMA strategy as well. The implementation of TCSS needs the ontology and annotation as text files provided by Gene Ontology Consortium. Therefore we use the released version of GO (gene_ontology.obo) dated Mar 13, 2015. The same released version of GO is used in above R packages (version: 3.1.2) and annotation for yeast (gene_association.sgd) and human (gene_association.goa_human) released on Mar 17,2015. We use ROC and ROCR R packages [92], [93] to plot the ROC curve and to calculate the area under ROC curves (AUC).

### 4.5 Results

Performance, in terms of Pearson correlation, of different SSMs with respect to the reference dataset from HIPPIE are shown in Table 3 (See Supplementary Material for barplots). The best correlations are shown in bold.

**TABLE 3.** Pearson Correlation with Reference Dataset from HIPPIE Database

| Ontology | Strategy | RDS | RNS | RES | TCSS | Resnik | Lin | Rel | Jiang | Wang |
| --- | --- | --- | --- | --- | --- | --- | --- | --- | --- | --- |
| BP | MAX | **0.358** | 0.313 | 0.346 | 0.342 | 0.329 | 0.277 | 0.277 | 0.272 | 0.286 |
|  | BMA | **0.342** | 0.332 | 0.310 | 0.270 | 0.238 | 0.220 | 0.218 | 0.211 | 0.193 |
| CC | MAX | 0.204 | 0.130 | 0.129 | **0.232** | 0.231 | 0.064 | 0.100 | 0.064 | 0.082 |
|  | BMA | **0.254** | 0.227 | 0.198 | 0.232 | 0.230 | 0.148 | 0.164 | 0.118 | 0.158 |

AUC obtained by different SSMs are tabulated in Table 4. The corresponding ROC curves are provided in the Supplementary Material. The best ROC scores are shown in bold.

**TABLE 4.**
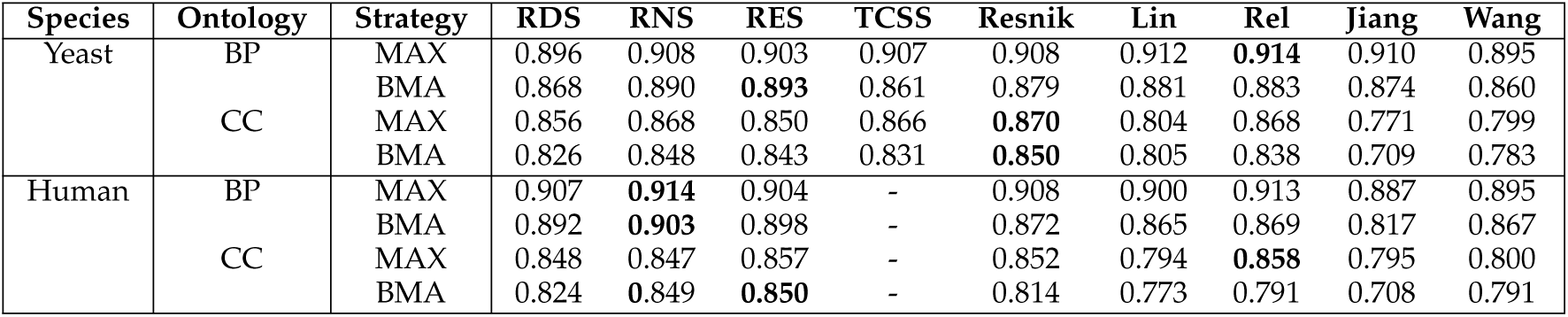
The area under ROC curves of different SSMs

As discussed earlier, the discriminating power quantifies the ability of an SSM to distinguish among various functionally different sets of proteins (e.g., KEGG pathways). Fig 1 and 2 demonstrate the discriminating power of different SSMs with BMA strategy against KEGG pathways in BP and CC ontology, respectively. Instead of pathway names, KEGG pathway identifiers are shown along the x-axis. The discriminating power for the selected yeast KEGG pathways (listed in Table 2) with more diverse functionality is shown in Fig 3. The results with MAX strategy are quite similar and kept in the Supplementary Material. The data tables are also provided in the Supplementary Material.

**Fig. 1.**
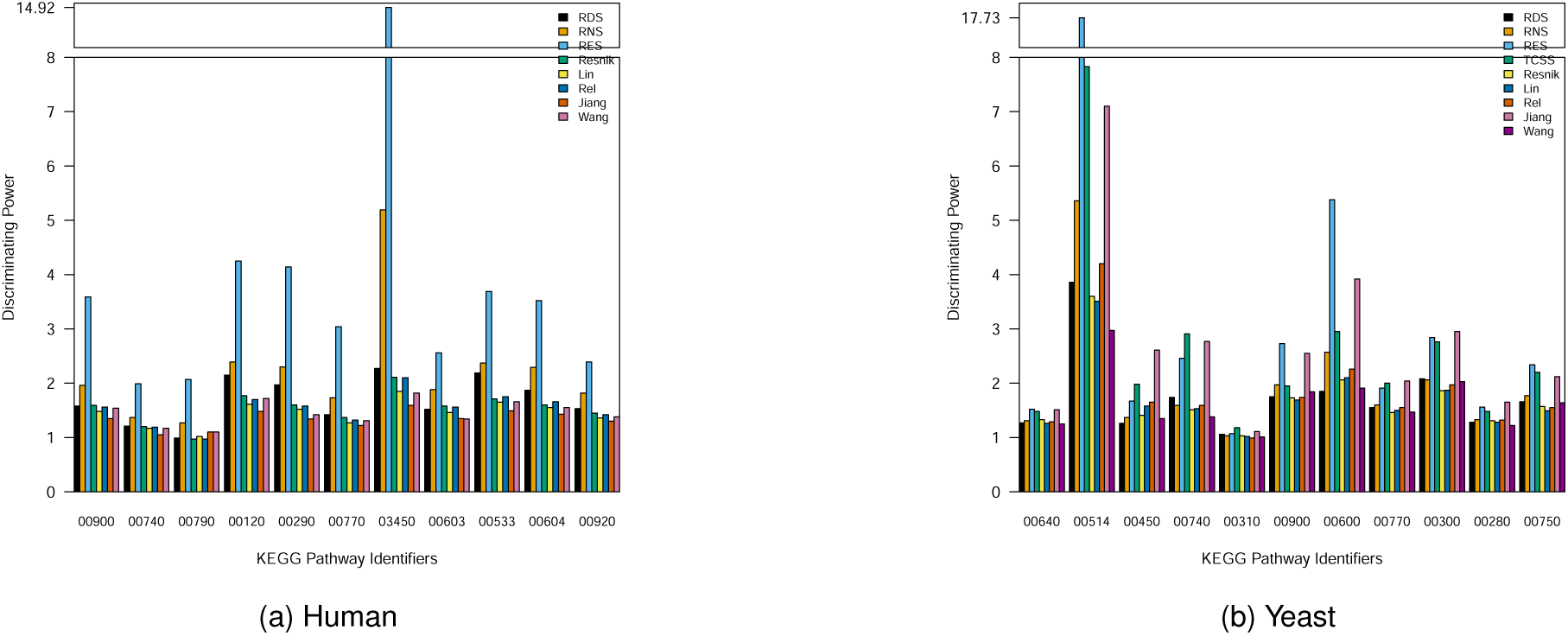
Inter-set discriminating power of different SSMs with BMA strategy in BP ontology. The y-axis is splitted to accommodate high DP value.

**Fig. 2.**
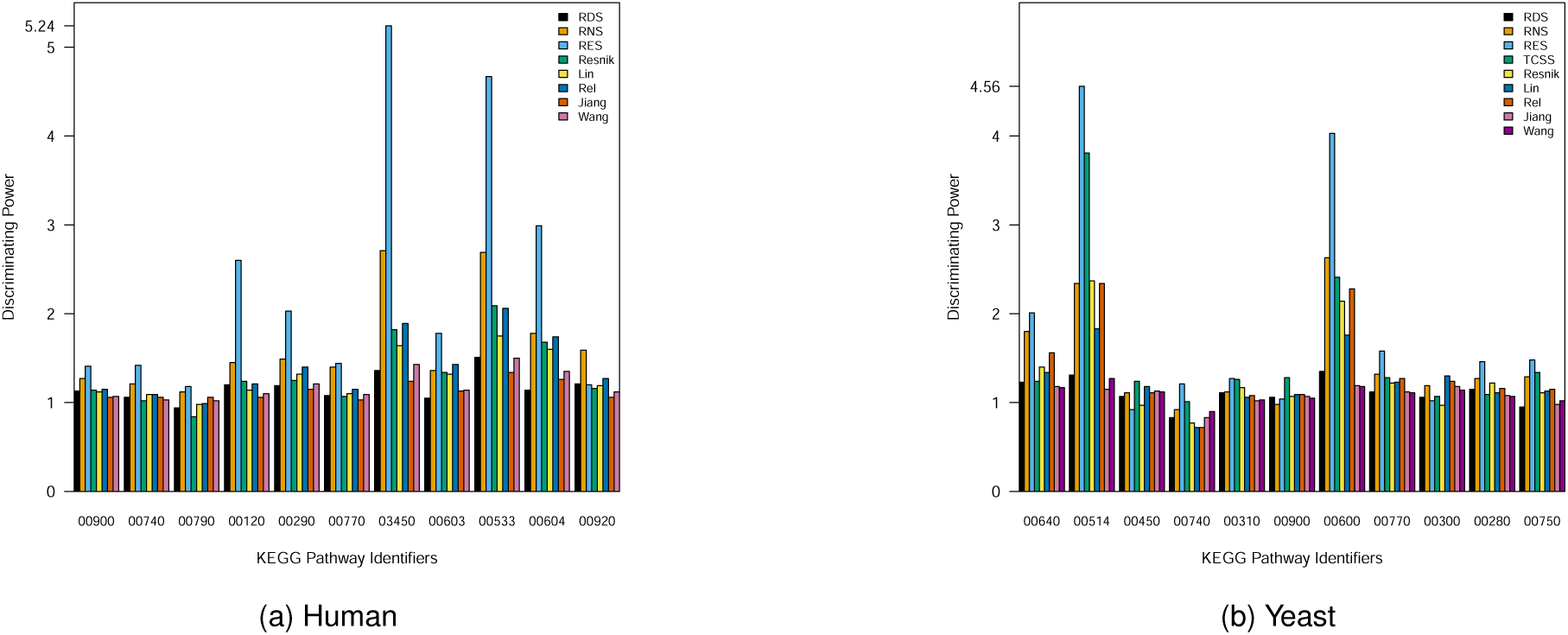
Inter-set discriminating power of different SSMs with BMA strategy in CC ontology.

**Fig. 3.**
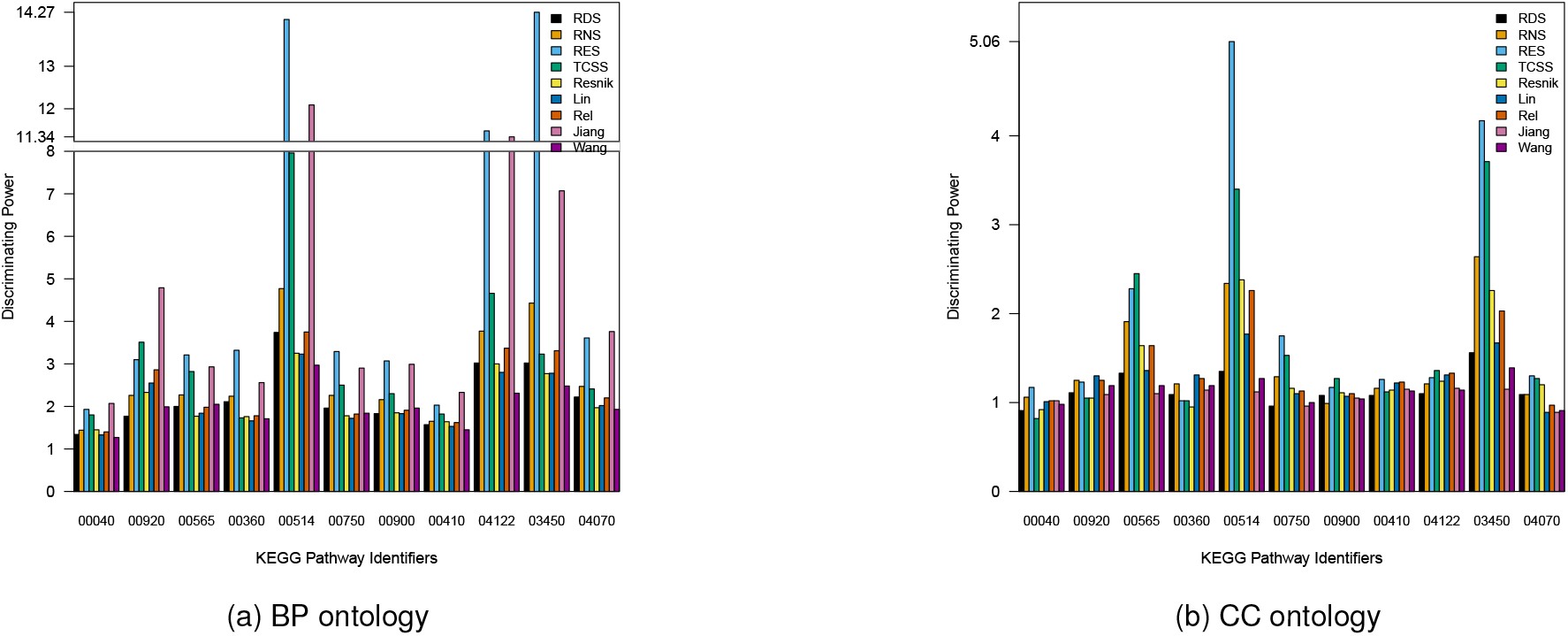
Inter-set discriminating power of different SSMs with BMA strategy for the selected 11 yeast KEGG pathways with more diverse functionality.

Finally, Table 5 demonstrates the performance of different SSMs on Pfam (See Supplementary Material for barplots). The best scores are shown in bold.

**TABLE 5.**
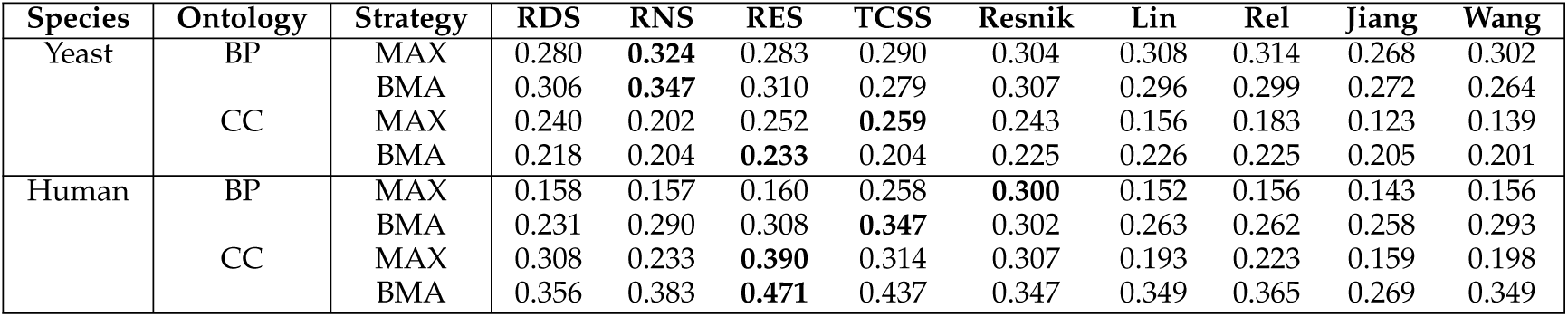
Correlation with Protein Family (Pfam) on CESSM dataset

## 5 Discussion

This section analyzes and discusses the results presented in section 4.5. We have highlighted the key observations.

### 5.1 Correlation with Reference Dataset from HIPPIE Database

#### RDS achieves the highest correlation in BP

while TCSS shows the maximum correlation in CC. It may be noted that RDS is the simplest SSM among the proposed measures and does not even consider annotation information. Nevertheless, it shows good correlation. RNS and RES also perform quite well in BP, while Resnik shows good performance in both BP and CC.

#### All SSMs show greater correlations in BP

The average correlation over all SSMs in BP is 0.311/0.259 (MAX/BMA), whereas in CC it is 0.137/0.192 (MAX/BMA). However, all measures show less overall correlation since correlation is computed for positive PPIs only.

### 5.2 ROC Curve Analysis

#### RNS and RES, with both MAX and BMA strategies, effectively classify true PPIs from false in both BP and CC

Resnik-MAX and Rel-MAX too perform well compared to others, while RDS shows competitive performance. Although we could not compare TCSS for human, it performs well with MAX strategy in yeast. All SSMs with MAX strategy have quite similar AUCs in BP for both yeast and human. However, with BMA strategy, AUCs achieved by RES (yeast:0.893, human:0.898) and RNS (yeast:0.890, human:0.903) are significantly higher than others. Further, RES and RNS exhibit greater consistency, since they show less difference between MAX and BMA strategies in both BP and CC (for both yeast and human).

#### All SSMs show higher AUCs in BP

The average AUCs in BP are 0.906/0.877 (yeast:MAX/BMA) and 0.904/0.873 (human:MAX/BMA), whereas in CC these are 0.839/0.815 (yeast:MAX/BMA) and 0.831/0.800 (human:MAX/BMA).

### 5.3 Set-discriminating Power of KEGG Pathways

**The discriminating power of RES is significantly higher** than other SSMs for all the 11 human KEGG pathways. RES produces DP value greater than or equal to 1.81/1.99 (MAX/BMA) in BP, while the next minimum DP value is 1.17 (produced by RDS - MAX).

#### RES shows maximum functional discrimination among the pathways

RES produces very high DP value with 11.10/14.92 (MAX/BMA) for *Non-homologous end-joining* (hsa03450) pathway. This is the only pathway that belongs to the *Genetic Information Processing* category, while rest fall in the same *Metabolism* category. So, the functional characteristic of *Non-homologous end-joining* pathway is completely different from the rest. RES nicely captures this functional discrimination by producing very high DP value.

#### All SSMs produce greater DP values in BP

Although RES almost consistently produces higher DP values in both BP and CC (with both MAX and BMA), it shows comparatively lower DP values in CC.

#### Overall discriminating power of all the SSMs are quite similar and not so good for the first set of yeast KEGG pathways

If we examine the functional categories of all the 11 pathways, we find that all belong to the same *Metabolism* category with six pathways from two subcategories only. Further, the selected first set of yeast pathways contain merely 134 genes with 16 are shared. In contrast, the selected human pathways include 150 genes with 11 are common only. Hence the selected first set of yeast pathways are functionally closer to each other and this fact is reflected by low DP values.

To study further, we consider another set of 11 yeast pathways with more diverse functionality, where three pathways (sec00514, sec00750, and sec00900) were taken from the previous set. The pathways are listed in Table 2 and corresponding discriminating power for BMA strategy is shown in Fig 3 (See Supplementary Material for data table).

#### The discriminating power of all the SSMs is improved significantly for the pathways with more diverse functionality

In particular, DP values of RES and Jiang are higher than other measures for almost all the pathways. RES and Jiang produce DP value greater than or equal to 2/1.93 (MAX/BMA) and 1.84/2.07 (MAX/BMA), respectively, in BP, while the next minimum DP value is 1.73 (produced by TCSS - BMA). The maximum DP value (MAX/BMA:13.75/14.27 in BP) is again produced by RES for the pathway sec03450 (Non-homologous end-joining).

#### RES can be used for functional clustering

It may be noted that although Jiang produces competitive DP values with RES for yeast pathways, it is unable to show good DP values for the human pathways. Therefore RES might be used for functional clustering (e.g., to characterize protein functional modules) as it shows consistently high discriminating power.

#### No SSM produces consistently good DP values in CC

particularly for the yeast pathways. Guo et al. [23] observed that all pairs of proteins involved in the same KEGG pathway have significantly higher similarity scores than randomly selected in BP, whereas similarity decreases exponentially as the distance between two proteins increases within the same pathway in CC and MF. These findings conform with current results as well.

### 5.4 Correlation with Pfam

#### Overall performances of TCSS, RES, and Resnik are well

Particularly, TCSS - MAX, RES - BMA, and Resnik - MAX perform well. Although RES does not show good correlation with MAX strategy in human, it produces a good correlation with BMA strategy. MAX strategy could overestimate while computing the general measure of functional similarity [22] and protein family captures a general aspect of protein function. Thus, BMA might be a better choice than MAX for Pfam similarity.

Further, it may be noted that correlation in CC is higher than BP in human for all measures which are quite unexpected. Therefore it might be challenging to draw comparative inference for the benchmark like Pfam that adopt a very general aspect of protein function with Jaccard index.

## 6 Conclusions and Future Work

The paper presents a new family of SSMs for scoring confidence of PPIs utilizing GO. This new family of SSMs is based on a new set of specificity measures namely, RDS, RNS and RES. Specificity of a term is redefined by considering the properties of its ancestors and descendants only along with its own properties so that maximum unwanted noises could be avoided. The evaluation shows that instead of simplicity, they are quite effective. Particularly, RNS and RES more effectively distinguish true interactions from false. RES can be useful for protein functional clustering as well since it shows a robust set-discriminating power over KEGG pathways. It also exhibits greater consistency and shows the best performance in BP with BMA strategy. Similar to the earlier studies, our evaluation also shows Resnik is one of the best SSMs for scoring confidence of PPIs. TCSS with MAX strategy and Rel also show competitive performance. Although RDS is the simplest SSM that does not even consider annotation information, it shows competitive performance as well. For almost all the four benchmarks, each SSM shows comparatively greater and consistent performances in BP. Therefore we believe that BP is more suitable than CC for scoring confidence of PPIs.

Although the newly developed SSMs are evaluated only on GO for scoring confidence of PPIs, it is not limited to any particular ontology. Therefore it would be worthy to evaluate how these SSMs perform on other ontologies and applications as future work.

## Availabilty of Data and Script

An R script for the new SSMs along with the complete datasets used in the evaluation is freely available at https://github.com/msp-cse/NaiveSSMs.

## Acknowledgments

The authors would like to thank Prof. V. Vijaya Saradhi for his valuable comments and suggestions.

---

**Madhusudan Paul** received the M.Tech degree in computer science and engineering from Pondicherry University in 2010. He is working towards the PhD degree in computer science and engineering from Indian Institute of Technology Guwahati, India. At present, he is an assistant professor at Visva-Bharati, Santiniketan, West Bengal, India. His current research interests include complex networks and systems biology.

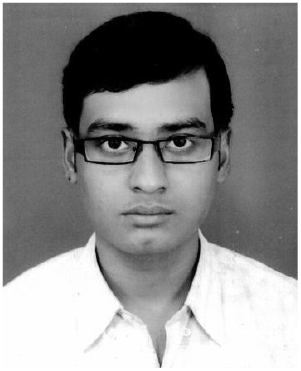

Ashish Anand is an associate professor at the Department of Computer Science and Engineering, Indian Institute of Technology Guwahati, India. His current research interests include Machine Learning and its application in computational biology, NLP, Clinical Text Mining, and Deep Learning. Web-site:

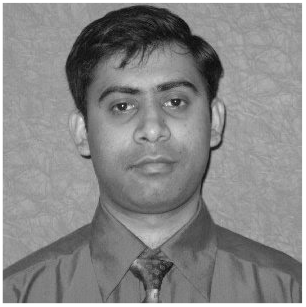

